# UPCLASS: a Deep Learning-based Classifier for UniProtKB Entry Publications

**DOI:** 10.1101/842062

**Authors:** Douglas Teodoro, Julien Knafou, Nona Naderi, Emilie Pasche, Julien Gobeill, Cecilia N. Arighi, Patrick Ruch

## Abstract

In the UniProt Knowledgebase (UniProtKB), publications providing evidence for a specific protein annotation entry are organized across different categories, such as function, interaction and expression, based on the type of data they contain. To provide a systematic way of categorizing computationally mapped bibliography in UniProt, we investigate a Convolution Neural Network (CNN) model to classify publications with accession annotations according to UniProtKB categories. The main challenge to categorize publications at the accession annotation level is that the same publication can be annotated with multiple proteins, and thus be associated to different category sets according to the evidence provided for the protein. We propose a model that divides the document into parts containing and not containing evidence for the protein annotation. Then, we use these parts to create different feature sets for each accession and feed them to separate layers of the network. The CNN model achieved a F1-score of 0.72, outperforming baseline models based on logistic regression and support vector machine by up to 22 and 18 percentage points, respectively. We believe that such approach could be used to systematically categorize the computationally mapped bibliography in UniProtKB, which represents a significant set of the publications, and help curators to decide whether a publication is relevant for further curation for a protein accession.

## Introduction

Due to the deluge of research data created at ever-increasing rates, the scientific community is shifting towards and relying on curated resources [1]. Biocurated resources provides scientists with structured, computable-form knowledge bases extracted from unstructured biological data, particularly published manuscripts, but also other sources, such as experimental data sets and unpublished data analysis results [2]. The UniProt Knowledgebase (UniProtKB) aims to collect functional information on proteins with accurate, consistent and rich annotation. In addition to capturing the amino acid sequence, protein name, taxonomic data and citation information, it includes literature-based information about different topics, such as function and subcellular location. UniProtKB combines reviewed UniProtKB/Swiss-Prot entries, to which data have been added by expert biocurators, with unreviewed UniProtKB/TrEMBL entries, which are annotated by automated systems, including rule-based [3]. The reviewed section represents less than 1% of the knowledgebase. Biocurators select a subset of the available literature for a given protein, representing the landscape of knowledge at a given time [4].

Given the extent of the datasets processed by biocurators, commonly in the range of thousands to millions of publications, the scalability of biocuration in life sciences has been often scrutinized [5].Text mining [6] has been proposed as one solution to scale up literature-based curation, especially to assist biocuration tasks via information retrieval, document triage, named entity recognition (NER) and relation extraction (RE), and resource categorization [7][8][9][10]. Textpresso Central, for instance, provides a curation framework powered with natural language processing (NLP) to support curators in search and annotation tasks in Wormbase and other databases [11]. Similarly, BioReader focus on the classification of candidate articles for triage [12]. Providing positive and negative examples, the framework is able to automatically select the best classifier among a range of classification algorithms, including support vector machine (SVM), k-nearest neighbours and decision tree. Tagtog leverages manual user annotation in combination with automatic machine-learned annotation to provide accurate identification of gene symbols and gene names in FlyBase [13]. Text mining has also supported more specific curation tasks, such as protein localization. LocText, for example, implements a NER and RE for proteins based on SVM, achieving 86% precision (56% F1-score) [14]. To address the common issue of class imbalance in biocuration, an ensemble of SVM classifiers along with random under-sampling were proposed for automatically identifying relevant papers for curation in the Gene Expression Database [15].

More recently, with the success of deep learning in image and text processing applications [16], deep learning models have been increasingly applied to biocuration. Deep learning classification and prediction models for text - the main use-cases in biocuration - are heavily supported by neural language models, such as word2vec [17] and Global Vectors (GloVe) [18], and lately by Embeddings from Language Models (ELMo) [19] and Bidirectional Encoder Representations from Transformers (BERT) [20]. Lee et al. used a convolutional neural network (CNN) model, supported by word2vec representations, in the triage phase of genomic variation resources, outperforming the precision of SVM models up to 3% [21]. This approach increased the precision by up to 1.8 times when compared to query-based triage methods of UniProtKB. Similarly, Burns et al. use a combination of CNN and recurrent neural network (RNN) models to scale-up the triage of molecular interaction publications [22].

To capture the breadth of publications about proteins and make it easily available to users, UniProt compiles additional bibliography from three types of external sources – databases, community and text mining – which complements the curated literature set with additional publications and adds relevant literature to entries not yet curated [4][23]. UniProtKB publications in reviewed entries are categorised on eleven predefined topics – *Expression, Family & Domains, Function, Interaction, Names, Pathology & Biotech, PTM / Processing, Sequences, Structure, Subcellular Location*, and *Miscellaneous* – based on the annotation they contribute to the protein entry, e.g., a paper that is the evidence source for the protein catalytic activity will be included in the category *Function* (Figure 1). On the other hand, to automatically categorize additional publications, UniProtKB uses the information from the underlying external sources (usually, databases). For example, the literature provided by the iPTMnet database is under the PTM / Processing category. Nevertheless, this approach is limited as it does not cover optimally all types of evidence available for the specific protein in the publication, other than PTM / Processing in this case. In fact, in publications imported from a number of sources, such as model organism databases, it is not possible to categorise unless the source provides the information.

**Figure 1.**
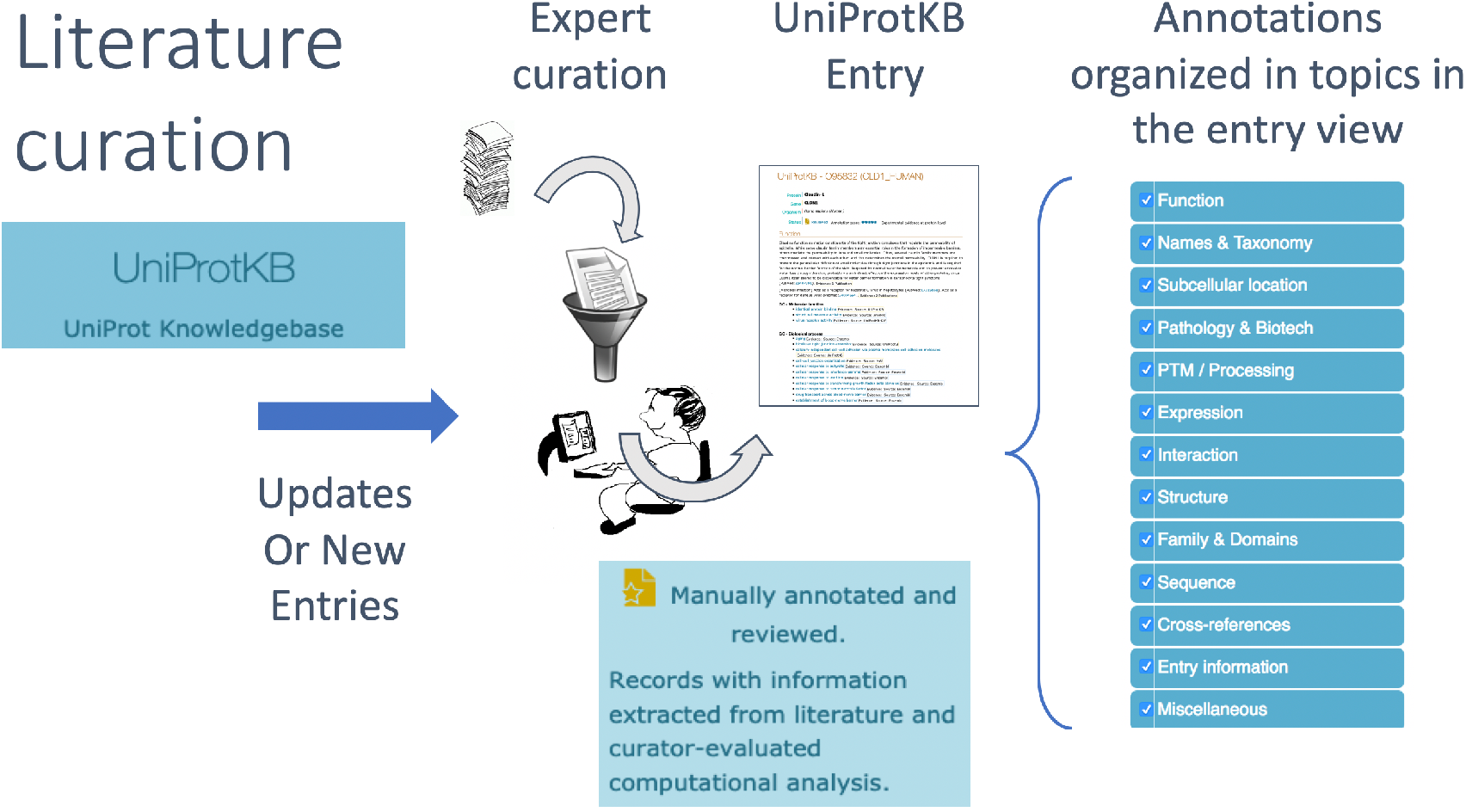
UniProt Knowledge annotation process. Manually protein-annotated documents (Swiss-Prot) are associated to UniProt Entry categories (Function, Name & Taxonomy, etc.) according to the type of information available in the publication, which are useful to organise the annotations within the knowledge base. A much larger set of publications (TrEMBL) is then automatically annotated according to their source characteristics.

Differently from classical text classification problems, it is often the case in biocuration where the same document can be associated to different class sets depending on the biological entity considered. For example, in UniProtKB, the same publication can be categorised into an entry set of the knowledge base for a protein *A* and into another entry set for protein *B* based on the evidence contained for each protein in the document. Standard document classification models cannot be generalised to this scenario as the input features are the same (i.e., the document) while the output classes are different (i.e., the entry set). To provide a systematic way of classifying the set of additional publications into the different UniProtKB annotation topics, in this paper we propose a model based on CNN to classify single publications into different class sets depending on the biological entity of interest. We use candidate evidence sentences, selected based on availability of protein information, to create different feature sets out of a unique document. These feature sets are then embedded into a continuous word representation space and used as input for a deep neural network-based classifier. We compare the effectiveness of the deep learning-based model with baseline classifiers based on logistic regression and SVM models.

## Materials and methods

To classify publications into the UniProtKB protein entry categories, we developed a text mining pipeline based on a CNN model, so called UPCLASS. The UPCLASS classification model was trained and evaluated using a large expert curated literature dataset available from UniProtKB. In this section, we describe the methods used to automatically and systematically classify scientific articles in the knowledge base.

### Candidate sentences for annotation evidence

A key challenge for classifying publications according to the UniProtKB entry categories is that, for the same document, a few to thousands of proteins can be annotated with different categories based on the evidence provided in the text. For example, as shown in Figure 2, if for protein *A* there are evidences in the article for the *Sequence* and *Function* categories, and for protein *B*, there is evidence only for the *Function* category, the publication will be annotated with different class sets for the different protein entries in knowledge base. However, since only a few articles are expert annotated per protein, usually with little redundancy on the type of information it brings to the knowledge base (i.e., an UniProtKB entry category), the protein itself cannot be directly used as a learning feature for a category because it is not an informative feature. In a classical document classification scenario, in which a label set is associated to a single document or to a document-accession pair, the classifier would always receive as input the same set of informative features (i.e., the document) independent of the labels associated to the pair document-accession. Thus, the classifier would not be able to learn the actual classes associated to the document-accession pair, as the triplet document-accession ➔ categories tends to appear only once in the knowledge base.

**Figure 2.**
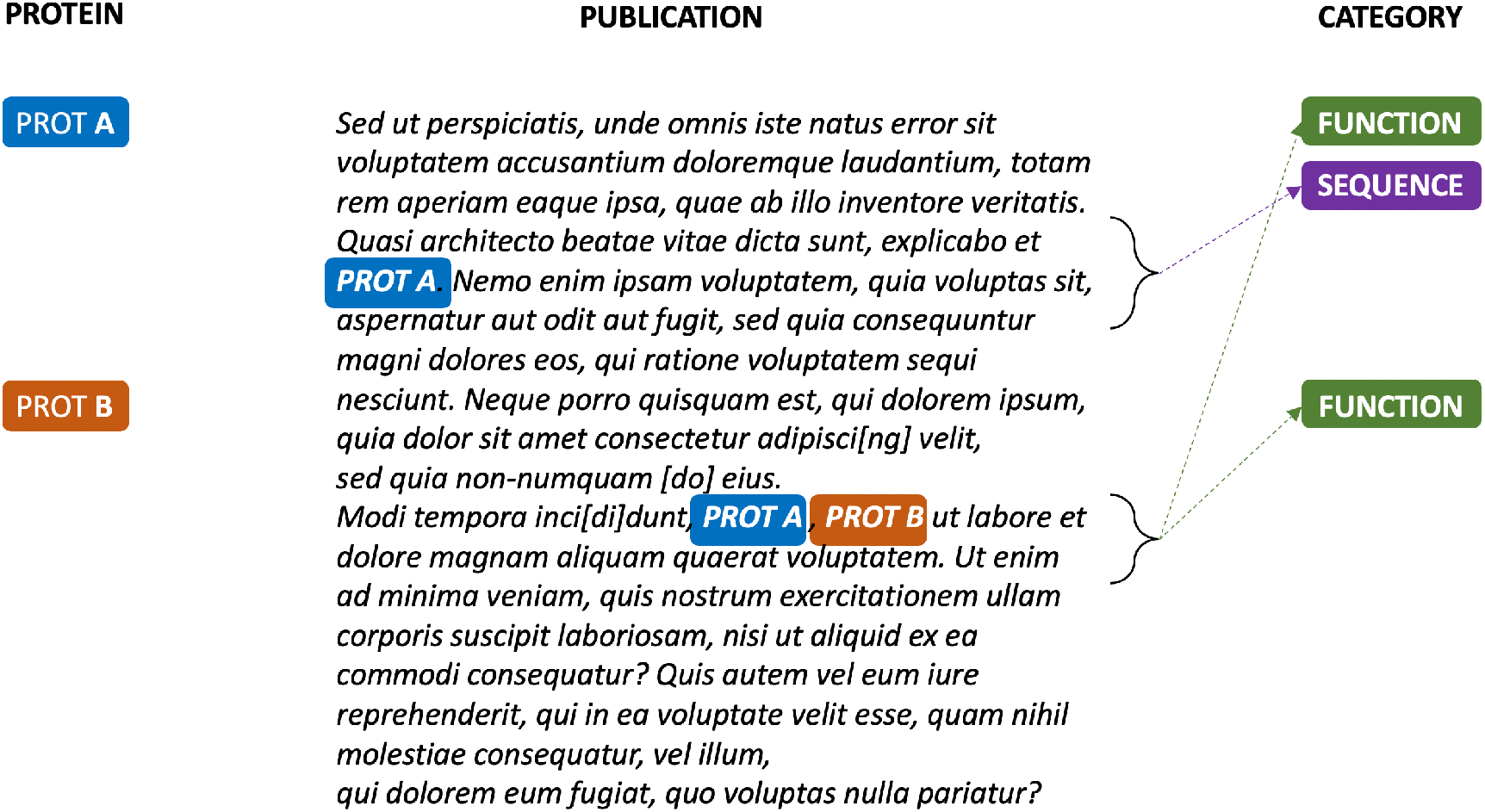
Synthetic annotation example illustrating how a single publication can be associated to different sets of UniProtKB entry categories.

To overcome this limitation, we developed a model where a document is dismembered into “positive” and “negative” sentences, based on whether they provide or do not provide evidences for the protein entry classification. Positive sentences are then concatenated to create a “positive” document for the respective protein annotation. The remaining sentences, i.e., the negative sentences, are similarly concatenated to create a “negative” document. Hence, as shown in Figure 3, different feature sets can be created out of a single document for each pair document-accession and be properly associated to their specific annotation categories.

**Figure 3.**
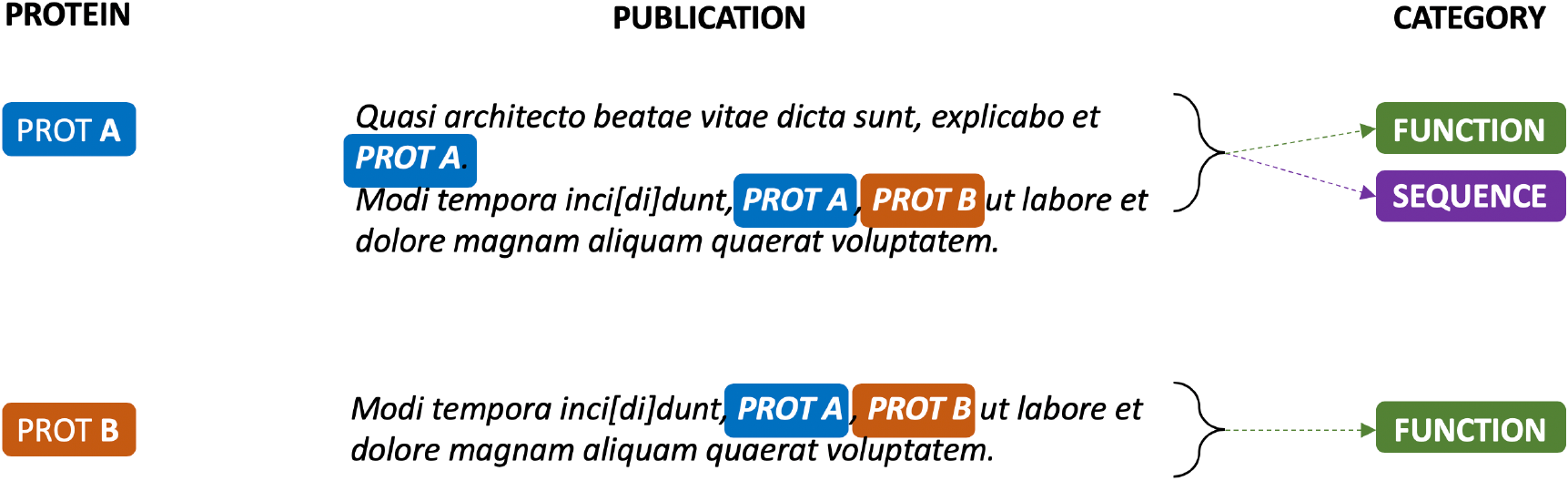
k-nearest (k=0) sentences containing candidate evidences for the protein annotation are concatenated to create a “positive” document. Similarly, sentences that do not contain evidence for the category are concatenated to create a “negative” document.

We hypothesize that the evidence for annotations is provided in the k-nearest sentences to the sentence where the protein (or its coding gene) is mentioned. Indeed, as shown by Cejuela et al. [14], the k-1 sentences accounts for 89% of all unique relationships in the case of protein location evidence. To identify the candidate evidence sentences, occurrence of protein features, such as accession identifier, protein name (recommended, alternative and short), gene name and their synonyms, are searched in the sentences. For example, for the accession number O95997, we search in the publication sentences for the strings “PTTG1_HUMAN” (accession), “Securin” (recommended name), “Esp1-associated protein” (alternative name), “Pituitary tumor-transforming gene 1 protein” (alternative name), “Tumor-transforming protein 1” (short name), “hPTTG” (short name), “PTTG1” (gene name), “EAP1” (gene synonym), “PTTG” (gene synonym) and “TUTR1” (gene synonym). If at least one match is found, the sentence is added to the positive pool. Subsequent sentences are further concatenated to form the “positive” document for the accession, O95997 in this case. These proteins features are available directly from the “Names & Taxonomy” section of the UniProtKB for each accession number.

### Classifier model

As illustrated in Figure 4, we use a three-layer CNN architecture for our machine learning classifier. The model has two branches, which receive the positive and negative documents separately. The architecture comprises three main building blocks: *i*) an input block (light blue), composed by an embedding layer (width=1500), which receives the document tokens and the pre-trained word vectors from a word2vec model trained on 10^6^ Medline abstracts with protein information; *ii*) a CNN block (orange), composed by three CNN layers with 128 channels, kernel width equal to 5, and ReLU activation, followed by a batch normalisation layer, which exposes a max pooling output followed by a drop-out layer (drop-out rate of 50%); and *iii*) a dense block (dark blue), composed by two dense layers, the first layer (width=128) receives the concatenated output of the CNN branches as input, followed by an output layer (width=11) with a softmax activation function. The outputs of the CNN branches are then concatenated and fed to the dense layer. The model was trained in 50 epochs with early stopping set for 5 consecutives epochs without improvement in the validation set. The model was implemented using the Keras framework in Python 3.

**Figure 4.**
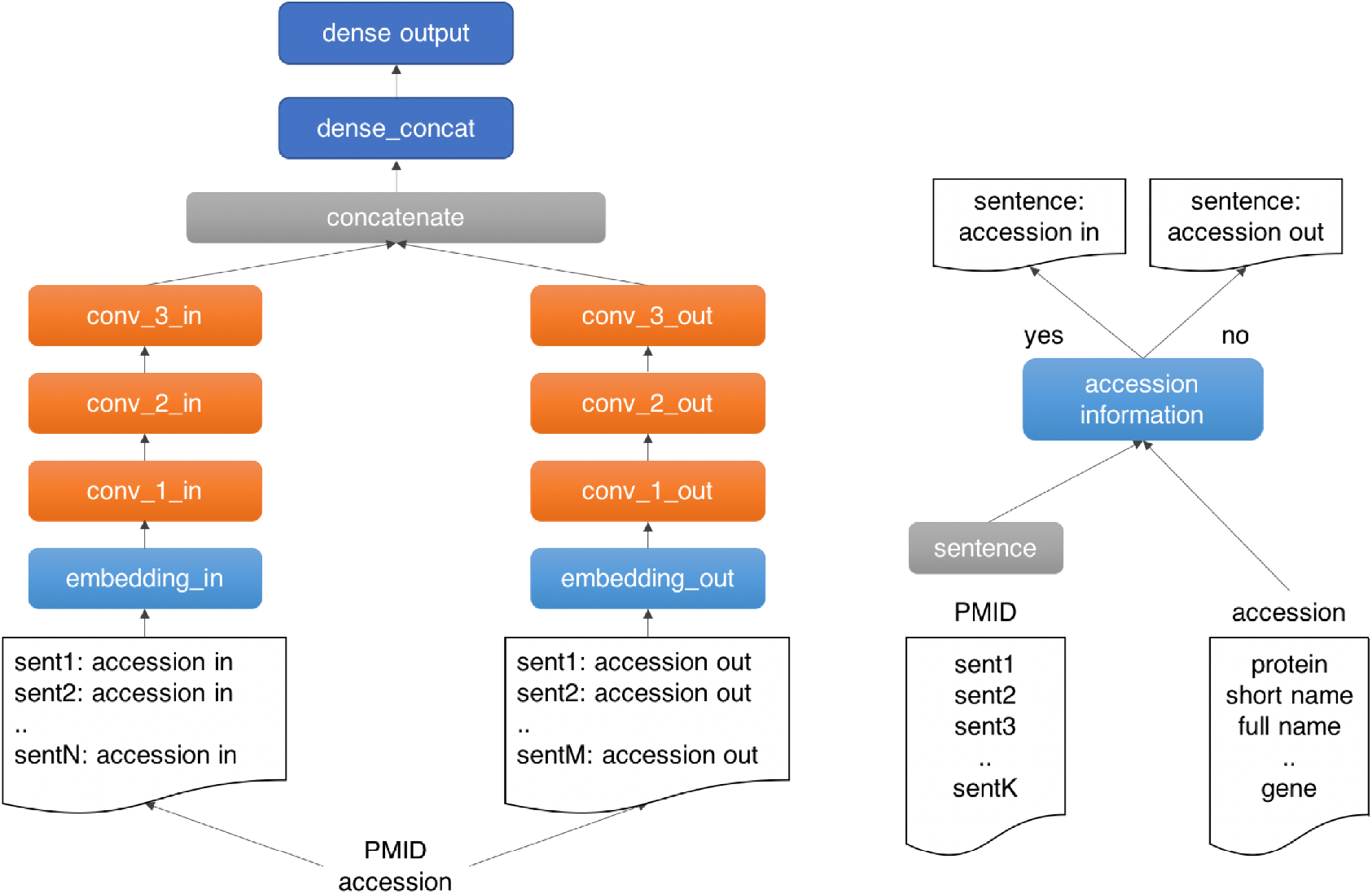
Outline of the UPCLASS CNN-based classification architecture with an embedding layer, three CNN layers followed by two dense layers. The “positive” sentences (accession in) are concatenated and fed to one branch of the model (“in” branch). The leftover sentences (accession out) are used to create the “negative” document and fed to the other branch of the model (“out” branch).

### Training and test collection

To train our model, we used an expert annotated collection of ~483k examples available from UniProtKB. In total, the collection contains ~201k unique manuscripts with an average of 2.4 proteins annotated per article (min=1, max=9329). The training collection was divided randomly in 76% for the train set (~368k samples), 12% for the dev set (~56k samples) and 12% for the test set (~58k samples). The division took into account the constraint that a publication should not be split in different sets, as the it is common the case when a protein is annotated for a category in a publication, other proteins share the same annotation. Full text publications were extracted from PubMed Central when available; otherwise, MEDLINE abstracts were used. In total, 96% of the collection was composed only by abstracts, with 5% of full text articles in the train set, 6% in the dev set and 8% in the test set.

In Table 1, the distribution of examples per category in the training collection is presented. As previously discussed, indeed the number of unique examples labelled per protein accession varies on average from 1.2 for the *Names* category to 2.2 for the *PTM / Processing* category. On the other hand, for categories like *Names*, *PTM / Processing* and *Sequences* there is less than one unique document per accession, i.e., it is often the case that a few proteins for these classes are annotated in the same publication. Moreover, we can notice that there is a concentration of samples in some classes, in particular *Sequences*, which is present in more than 45% of the examples, while *Structure*, *Names* and *Family & Domains* are present in 5% or less.

**Table 1.** Distribution of categories in the manually annotated training collection from UniProtKB. Examples: number of document-accession examples annotated with a category in the training collection. Unique accessions: number of unique accessions annotated with a category in the training set. Unique document: number of unique publications annotated with a category in the training set.

| UniProtKB entry category | Examples | Unique accessions | Unique documents |
| --- | --- | --- | --- |
| Expression | 53274 | 35128 | 34446 |
| Family & Domains | 4910 | 3807 | 3310 |
| Function | 105417 | 49896 | 72674 |
| Interaction | 60252 | 28318 | 30646 |
| Names | 11334 | 9130 | 1100 |
| Pathology & Biotech | 39870 | 23573 | 32410 |
| PTM / Processing | 69080 | 31142 | 17335 |
| Sequences | 217879 | 130288 | 109333 |
| Structure | 25569 | 14257 | 19553 |
| Subcellular Location | 48876 | 31866 | 28793 |
| Miscellaneous | 111454 | 52724 | 16812 |
| **Total** | **483159** | **163913** | **201358** |

### Pre-processing and word embeddings

In the pre-processing phase, we treat the training collection and the protein features (name, gene, etc.) through an NLP pipeline. First sentences are split using a Punkt sentence tokenizer. Then, stopwords are removed, characters are converted to lower case and non-alphanumerical characters are suppressed. The resulting tokens are stemmed and stem words smaller than 2 characters are removed. Finally, numerical sequences are replaced by the token *_NUMBER_*. Table 2 shows an example of the resulting sentences after passing the manuscript through the pre-processing pipeline.

**Table 2.** Resulting sentences after passing through the pre-processing pipeline.

| Original text | Pre-processed sentences |
| --- | --- |
| YddV from Escherichia coli (Ec) is a novel globin-coupled heme-based oxygen sensor protein displaying diguanylate cyclase activity in response to oxygen availability. In this study, we quantified the turnover numbers of the active [Fe(III), 0.066 min(-1); Fe(II)-O(2) and Fe(II)-CO, 0.022 min(-1)] [Fe(III), Fe(III)-protoporphyrin IX complex; Fe(II), Fe(II)-protoporphyrin IX complex] and inactive forms [Fe(II) and Fe(II)-NO, <0.01 min(-1)] of YddV for the first time. | yddv escherichia coli ec novel globin coupl heme base oxygen sensor protein display diguanyl cyclas activ respons oxygen avail<br>studi quantifi turnov number activ fe iii<br>_NUMBER_ min fe ii fe ii co<br>_NUMBER_ min fe iii fe iii<br>protoporphyrin ix complex fe ii fe ii<br>protoporphyrin ix complex inact form fe ii fe ii _NUMBER_ min yddv first time |

The word embedding weights were pre-trained in a pararaph2vec to model using the train set collection [24]. The motivation was that, in addition to adapt to the preprocessed text, as showed by Diaz *et al.*, locally trained word embeddings provide superior word representation [25]. Two paragraph2vec models were trained – DBOW (Distributed Bag of Words) and DMC (Distributed Memory Concatenated) – through 100 epochs with word and document vector size of 200 and window of 10, the optimal values found during the training phase for the classification models. The gensim Python library was used to the train the pararaph2vec models.

### Evaluation criteria

Results are reported using standard multi-label classification metrics - Precision, Recall and F1-score - and are compared to a baseline model based on logistic regression. Student’s t-test are used to compare the classifier models and results are deemed statistically significant for p-value < .05. As most of the publications contain more than one annotation per protein and they are often classified into the same classes, e.g., a paper containing structure information for several proteins, the predictions might not be independent for each sample. Thus, we report results for real curation use-cases but also considering only one unique random annotation per publication. Finally, it is also important to notice that a system that provides automatic annotations to knowledge bases should aim first at high precision. Nevertheless, in our case, we expect the classifier to go beyond the coverage provided by standard provenance classification, and, hence, demonstrate also high recall.

## Results

Table 3 shows the performance of the classification models used to categorize document-protein accession pairs according to UniProtKB entries. Three classification models were assessed -logistic regression (baseline), SVM and CNN - in two versions: “not tagged” and “tagged”. The “not tagged” version of the logistic regression and SVM classifiers received as input a 400-dimensional feature vector created from the concatenated output of the DBOW and DMC paragraph2vec models applied to a preprocessed publication. The “tagged” version received as input an 800-dimensional feature vector, 400 for the positive document and 400 for the negative document, created by tagging protein features against the publication sentences. Similarly, the CNN “not tagged” model received a 1500 token vector per document in the embedding layer while the CNN “tagged” model received a 1500 token vector for each branch of the model (positive and negative). The token weights for the embedding layer of the CNN models were provided by the trained DBOW document embedding model.

**Table 3.** Micro and macro average results for the *not tagged* and *tagged* models obtained from the test set of 58k records. Highest results are showed in bold. * statistically significant improvement.

| Model | Micro |  |  | Macro |  |  |
| --- | --- | --- | --- | --- | --- | --- |
|  | Precision | Recall | F1-score | Precision | Recall | F1-score |
| Logistic not tagged | 0.63 | 0.42 | 0.50 | 0.55 | 0.42 | 0.50 |
| Logistic tagged | 0.55 | 0.66 | 0.60 | 0.48 | 0.60 | 0.53 |
| SVM not tagged | 0.74 | 0.43 | 0.54 | 0.56 | 0.28 | 0.37 |
| SVM tagged | **0.75\*** | 0.38 | 0.50 | 0.64 | 0.25 | 0.36 |
| CNN not tagged | 0.67 | **0.76\*** | 0.71 | **0.68\*** | 0.46 | 0.55 |
| CNN tagged | 0.69 | 0.74 | **0.72\*** | 0.60 | **0.63\*** | **0.62\*** |

Overall, the CNN *tagged* model achieved the highest performance in terms of the F1-score metrics, outperforming all the other models for both micro and macro averages (p < .05). It outperformed the baseline logistic regression classifier in absolute values by 12% and by 9% for the micro and macro F1-score metrics, respectively. The SVM tagged *model* achieved the highest micro precision and the CNN *not tagged* model achieved the highest macro precision (p < .05), both at the expense of recall. Similarly, recall performance varies depending on how the results are aggregated. Micro recall is highest for the CNN *not tagged* model and macro recall is highest for the CNN *tagged* model. Since some of the categories had relatively few examples in the training set, the macro average metrics provide better insights on how the models are able to deal with class imbalance, as the macro metrics treat all classes equally, independent of their frequency in the training set. To this end, the CNN *tagged* model has an outstanding performance, increasing the F1-score metric by 7% when compared to the CNN *not tagged* model.

As publications are often annotated for several proteins (thousands in some cases), the pairs document-accession are not necessarily independent samples. Thus, we modified the test set to contain only one sample of a document per unique category set. Accession with the same annotation for a publication were randomly suppressed from the collection. This resulted in a test set of ~26k records, a reduction of 55% when compared to the original set of ~58k samples. The results of these tests are shown in Table 4. There was a relevant drop in performance for the CNN models, e.g., 4% and 6% in micro average F1-score for the *not tagged* and *tagged* models, respectively, and an overall relative improvement in relation to the CNN models for the logistic and SVM models. Nevertheless, the CNN models still outperform the baseline and SVM when considered the F1-score metrics. The best model in this setting is now the CNN *not tagged*, with F1-scores of 0.67 and 0.54 for the micro and macro averages, respectively. Thus, for an individual independent classification, the CNN *not tagged* classifier is likely to provide the best answer while for a collection with the similar distribution to UniProtKB’s categories, the CNN *tagged* model provides the best results.

**Table 4.** Micro and macro average results for the *not tagged* and *tagged* models obtained from the test set of unique document ➔ categories pairs (around 26k samples). Highest results are showed in bold. * statistically significant improvement.

| Model | Micro |  |  | Macro |  |  |
| --- | --- | --- | --- | --- | --- | --- |
|  | Precision | Recall | F1-score | Precision | Recall | F1-score |
| **Logistic not tagged** | 0.56 | 0.68 | 0.61 | 0.45 | **0.56\*** | 0.50 |
| **Logistic tagged** | 0.59 | 0.66 | 0.62 | 0.46 | 0.53 | 0.49 |
| **SVM not tagged** | 0.73 | 0.43 | 0.54 | **0.62\*** | 0.25 | 0.36 |
| **SVM tagged** | **0.75\*** | 0.45 | 0.56 | 0.61 | 0.27 | 0.38 |
| **CNN not tagged** | 0.64 | **0.71\*** | **0.67\*** | 0.54 | 0.54 | **0.54\*** |
| **CNN tagged** | 0.65 | 0.66 | 0.66 | 0.55 | 0.49 | 0.52 |

### Prediction comparison for the not tagged and tagged models

Out of the ~22k unique publications in the test set, around 7k (31%) were labelled with two or more category sets. In Table 5, we show three non-exhaustive examples, highlighting the different outcomes for the *not tagged* and *tagged* CNN classifiers for such cases. For the first publication, PMID 11847227, five distinct category sets were annotated for to the different accessions. As expected, the *not tagged* model associated only one type of category set to all document-accession pairs while the *tagged* model changed the predicted classes according to the accession features. This led to an increase in the micro average F1-score from 0.37 for the *not tagged* to 0.73 for the *tagged* results. On the second example, PMID 15326186, both models behave similarly, providing only one set of categories, independent of the accession information. This situation is usually seen for the *tagged* model when the classifier is not able to tag accession information in the document or all sentences in the abstract contains a protein feature token. Thus, a unique set of features, i.e., the publication vector, is used for classification. Finally, on the third example, PMID 10427773, the *tagged* model has more false positive predictions, lowering its precision when compared to the *not tagged* model by 2% (F1-score of 0.83 and 0.81, respectively).

**Table 5.**
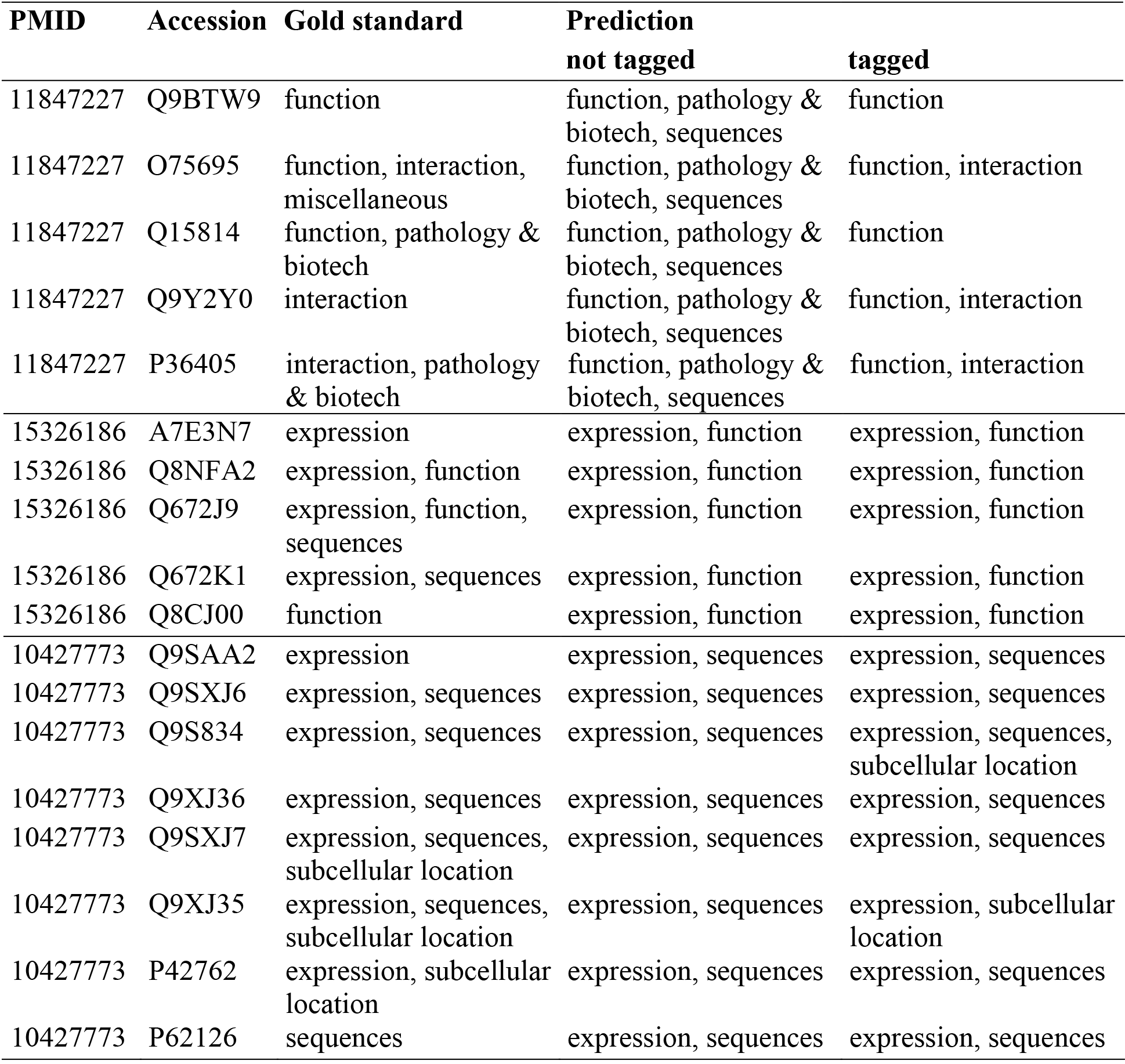
Examples of prediction output for the CNN *not tagged* and *tagged* models. While the CNN *not tagged* model provides only one type of output independent of the accession evidence in the publication, the *tagged* model attempts to predict the categories according to the accession features. For brevity, only unique examples of document ➔ categories pairs are shown.

| PMID | Accession | Gold standard | Prediction |  |
| --- | --- | --- | --- | --- |
|  |  |  | not tagged | tagged |
| 11847227 | Q9BTW9 | function | function, pathology & biotech, sequences | function |
| 11847227 | O75695 | function, interaction, miscellaneous | function, pathology & biotech, sequences | function, interaction |
| 11847227 | Q15814 | function, pathology & biotech | function, pathology & biotech, sequences | function |
| 11847227 | Q9Y2Y0 | interaction | function, pathology & biotech, sequences | function, interaction |
| 11847227 | P36405 | interaction, pathology & biotech | function, pathology & biotech, sequences | function, interaction |
| 15326186 | A7E3N7 | expression | expression, function | expression, function |
| 15326186 | Q8NFA2 | expression, function | expression, function | expression, function |
| 15326186 | Q672J9 | expression, function, sequences | expression, function | expression, function |
| 15326186 | Q672K1 | expression, sequences | expression, function | expression, function |
| 15326186 | Q8CJ00 | function | expression, function | expression, function |
| 10427773 | Q9SAA2 | expression | expression, sequences | expression, sequences |
| 10427773 | Q9SXJ6 | expression, sequences | expression, sequences | expression, sequences |
| 10427773 | Q9S834 | expression, sequences | expression, sequences | expression, sequences, subcellular location |
| 10427773 | Q9XJ36 | expression, sequences | expression, sequences | expression, sequences |
| 10427773 | Q9SXJ7 | expression, sequences, subcellular location | expression, sequences | expression, sequences |
| 10427773 | Q9XJ35 | expression, sequences, subcellular location | expression, sequences | expression, subcellular location |
| 10427773 | P42762 | expression, subcellular location | expression, sequences | expression, sequences |
| 10427773 | P62126 | sequences | expression, sequences | expression, sequences |

### Classifier performance analyses

As showed in Figure 5, to understand the impact of the different categories in the classifier performance, we analysed the precision-recall curve for the CNN *tagged* model (other classifiers show similar curves). Three categories have mean average precision (MAP) above 75%: *Sequences*, *Structure* and *Miscellaneous*. On the other hand, three categories have MAP lower than 50%: *Expression*, *Family & Domains* and *Pathology & Biotech*. The correlation between the number of examples in the training set and the category performance is only moderate (r = 0.53). Indeed, the *Structure* category, for example, is only present in 5% of the training examples, however it has one of the highest MAP. Differently, the *Expression* category is present in 11% of the examples (the median value among the categories) but has one of the lowest MAP. The *Family & Domains* category, which has only 1% of the training set records, was not learned by the classifier (MAP = 0.07). However, the *Names* category has 2% of the examples and a MAP ten times higher (MAP = 0.73). Hence, the class imbalance alone is not enough to explain the difference in performance among the categories.

**Figure 5.**
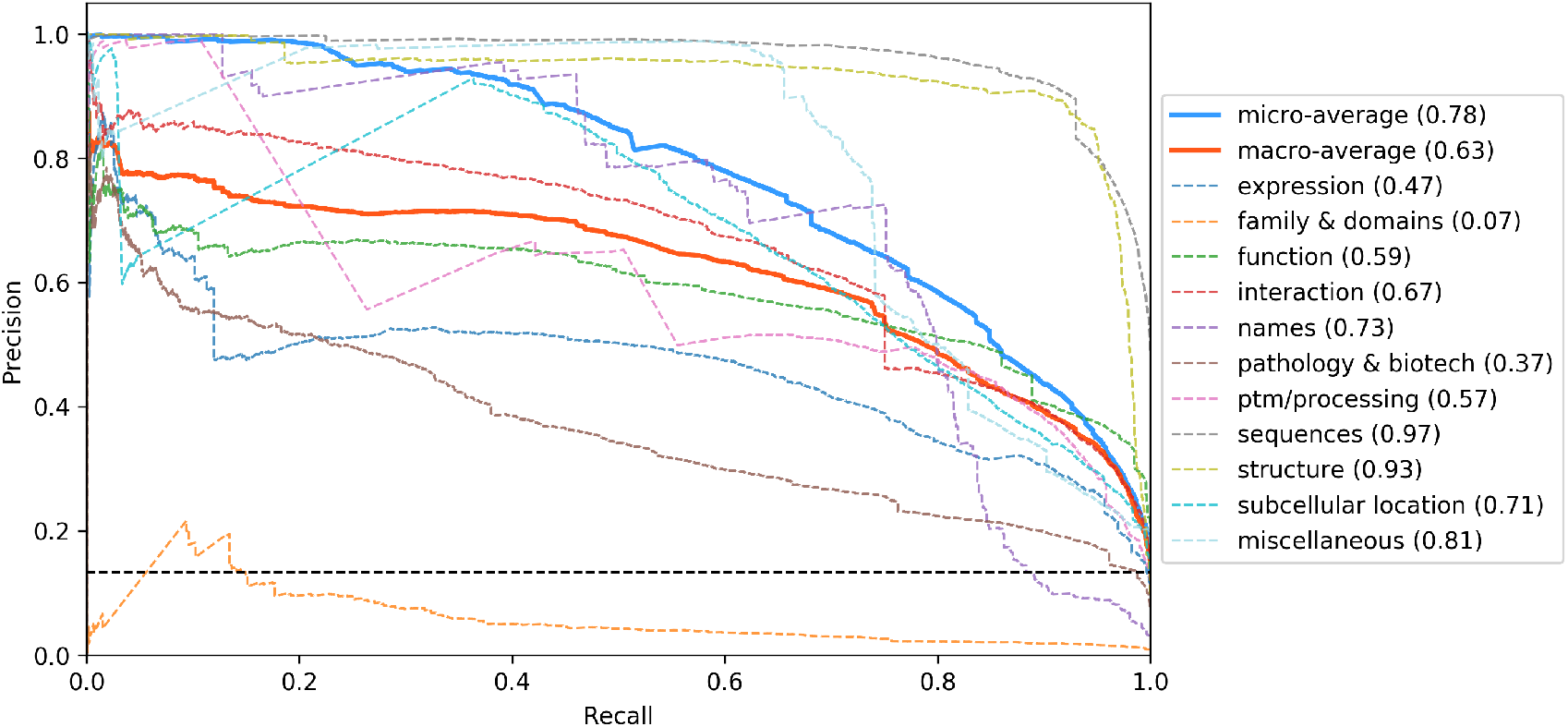
Precision-recall curves for the UniProtKB categories obtained from the CNN tagged classification. Mean average precision is shown in parentheses. Black horizontal dashed line: performance of a random classifier.

In Figure 6, the classifier prediction as a result of the input size is shown. Documents with size around 4k bytes contained only the abstract section and composed more than 95% of the test set. The remaining 5% were composed by full text documents or with at least some extra sections in addition to abstract, as figure and table captions. The correlation of the classifier performance and the size of the publication (or the type of annotated publication: abstract or full text) is very weak (*ρ* = 0.09). Thus, it seems that most of the evidence for the categories are provided in the abstract, apart from the *Expression*, *Family & Domains* and *Pathology & Biotech* categories, which have a MAP lower than 0.5.

**Figure 6.**
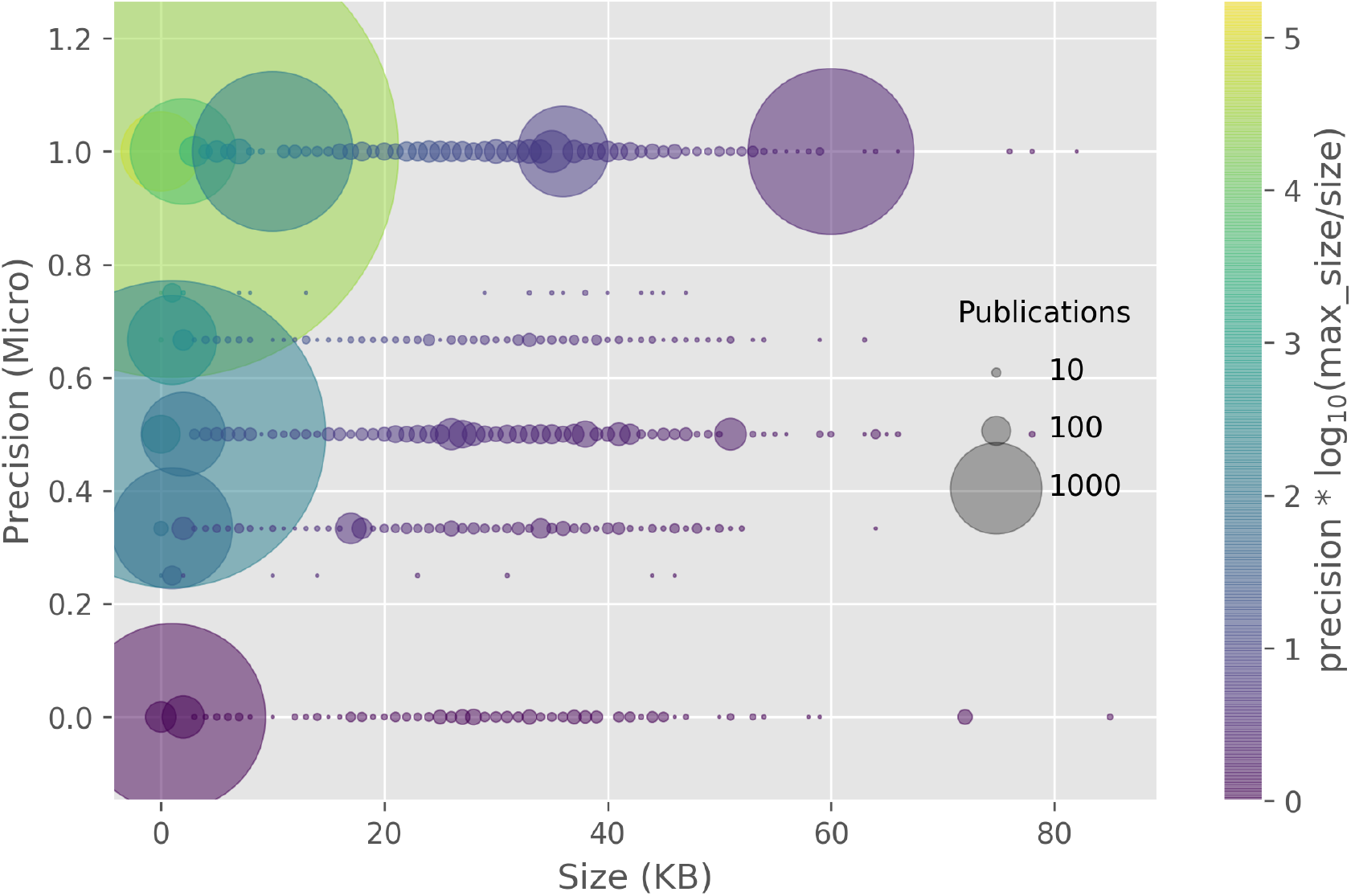
Classifier precision as a function of the publication size. There is no correlation between the size of the input size and precision. Circle size: number of publications within a size bin. Yellow points: high precision and lower ratio between publication size and the max publication size in the test set. Purple points: low precision and higher ratio between the publication size and the max publication size in the test set.

Finally, on Figure 7 we show the micro average precision results of the CNN *tagged* classifier for the 19 most common organisms in the test set. For the organisms with higher precision, *Drosophila melanogaster* (DROME), *Schizosaccharomyces pombe* (SCHPO), and *Oryza sativa subsp. Japonica* (ORYSJ), the majority of the protein entry annotations belong to a few classes (median frequency of classes of less than 2%). These organisms were mostly annotated for *Sequences* and *Subcellular Location* categories (~60% or more of the annotations), which had overall high to moderate-to-high precision performance, respectively, as shown in Figure 5. Conversely, organisms with well distributed annotations among the 11 categories in the test set (median distribution of ~9%) had the poorest precision scores: *Arabidopsis thaliana* (ARATH), *Candida albicans* (CANAL), and *Dictyostelium discoideum* (DICDI). The low performing categories are likely to have impacted negatively the precision of these organisms.

**Figure 7.**
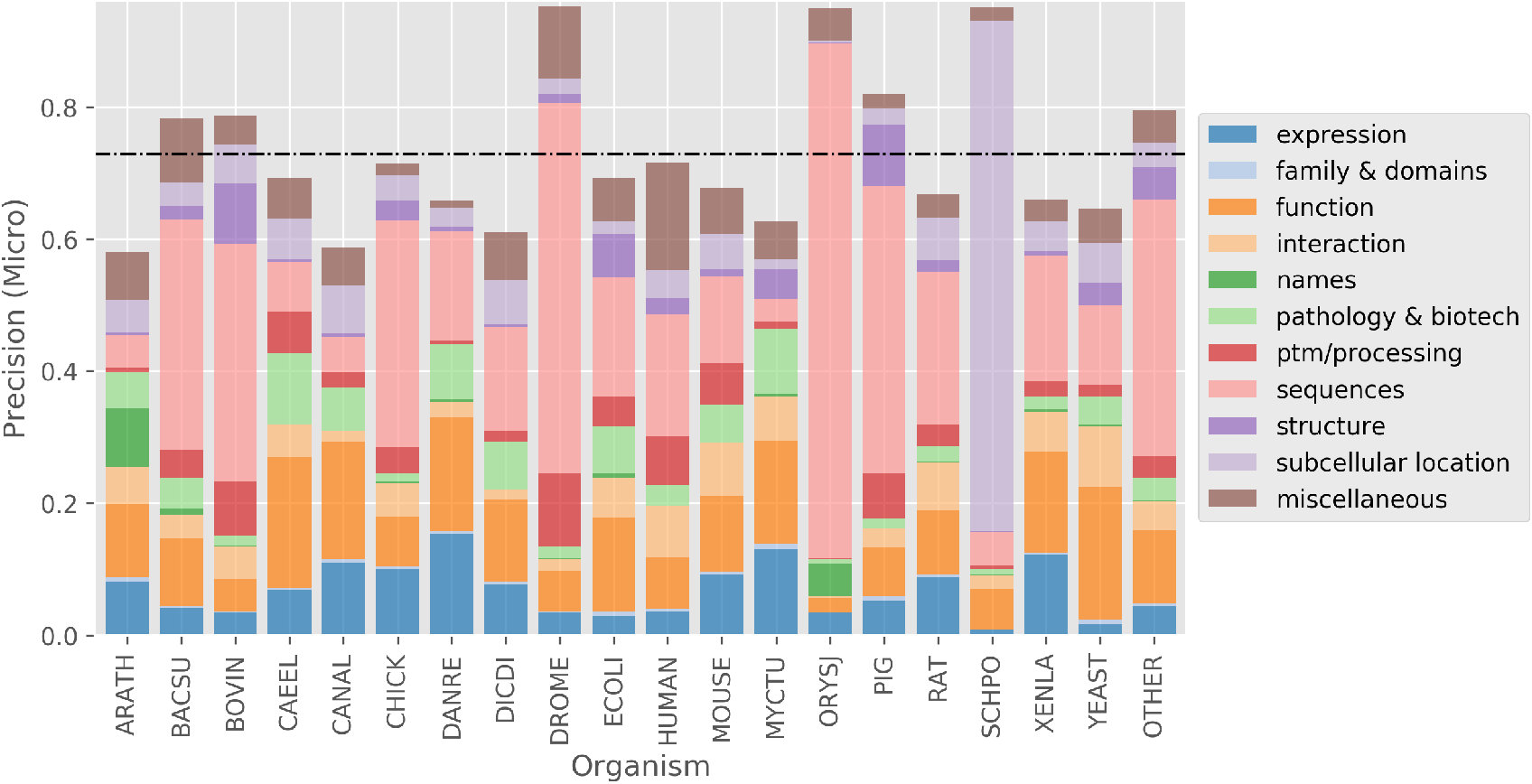
Micro average precision performance per organism. Higher precision for organisms happens when there is a concentration of categories. Black dashed horizontal line: mean organism precision.

### UniProtKB category evidence annotation

To measure the performance of the tagging method for detecting candidate evidence sentences, we compared sentences from 20 manually annotated abstracts with the positive and negative sentences created by our naïve string-matching algorithm. In total, 61 sentences distributed among 7 categories - *Expression, Function, Interaction, Pathology & Biotech, PTM / Processing, Sequence* and *Subcellular Location* - were tagged as positive. The algorithm achieved micro precision of 0.42 (28 out of 67 sentences) and micro recall of 0.46 (28 out of 61 sentences). For 4 abstracts (20%), no positive sentence was detected. In this case, the whole abstract was considered as positive. For the other 5 articles (25%), no true positive sentence was tagged. After assessing the classification performance for this test set (F1-score of 0.70), there was no correlation between the detection of candidate sentences and the prediction of the individual document-accession pairs (ρ = 0.02).

## Discussions

We investigated the use of a supervised CNN classifier to automatically assign categories to document-accession pairs curated in the UniProtKB to help scaling up publication categorization into the knowledge base entry categories. The classifier was trained and evaluated using a collection of 483k document-accession pairs annotated biocurator experts. To overcome the issue of multiple category sets associated to a single publication, we proposed an effective strategy to tag publication sentences with protein features and create different feature sets out of a single document entry. Results showed statistically significant improvements upon models that use only the publication as classification features, improving F1-score up to 12% (micro) when compared to the logistic regression baseline and up to 7% (macro) when compared to the *not tagged* CNN version.

While for some database resources it has been shown that expert curation can keep up with the exponential growth of the scientific literature [4], scaling-up biocuration remains a challenge. A key success factor for UniProtKB’s scalability, for example, is that the set of expert curated literature in the knowledge base focus on non-redundant annotations for proteins. It relies on external sources for the contribution of additional literature and on UPCLASS for its classification. Indeed, the implementation of UPCLASS has enabled the classification of more than 30 million document-accession pairs according to the entry categories, which were previously displayed as unclassified in UniProtKB [3]. UPCLASS is publicly available as a web service through the URL address: http://goldorak.hesge.ch/bioexpclass/upclass/.

In addition to direct classification, models as provided by UPCLASS could be used in other automatic phases of the biocuration process, such as document triage, helping curators to reduce the search scope. In the context of UniProtKB curation workflow, it could be used to prioritize UniProtKB entries that are unreviewed proteins with publications in the additional bibliography belonging to some category of interest, or to update reviewed entries that lack *Function* annotation, for example, but for which there are papers in the additional bibliography section classified by UPCLASS in the *Function* category. In [10], UPCLASS was used in a question-answer model to classify the type of search questions and their respective result sets in biomedical metadata repositories, e.g., search for gene expression data or search for protein sequencing data. While this approach did not lead to improvements in information retrieval performance of datasets, further investigation is needed, in particular in a context where the type of classification material is aligned with the training set type. Overall, given the outstanding results provided by the CNN models when compared to the baseline model but even to more powerful frameworks, such as SVM, we expect that such models could be extended to other literature curation domains, for example, in prior art search and classification of patents [26][27].

### Classification performance per category, document size and organism

We analysed the performance of the CNN *tagged* classifier according to three dimensions: category, document size and annotated organism. Despite the good performance of the classification models, there is still room for improving precision for some of the UniProtKB classes. Several attempts were made to increase UPCLASS outcome using imbalanced learning methods, such as under sampling and Tomek links [28]. However, they did not lead to overall improvement. An alternative would be to use a classifier ensemble, as proposed in [15]. However, this approach is too expensive for deep learning models due to the learning cost. We also tried to change the number of k-nearest sentences in the tagged models, but it did not lead to positive changes in performance either. It would be interesting to explore the addition of expert categorisation rules, in particular of UniProtKB mapping rules, as a strategy to increase performance while keeping the training complexity relatively low. The analyses of the classification precision according to the size (or type) of the document shows that there is no correlation between these two dimensions. While it might be counter-intuitive, several works have demonstrated that for classification tasks, the performance of abstracts are at least equivalent to full texts, when not better [29]. Yet, it could be relevant to explore the use of different classifiers for abstracts and for full texts. Finally, performance analyses according to annotated organism shows that better prediction outcome are mostly a result of high-performing class annotations being concentrated on some organisms. Thus, it seems that the classification precision for organisms is not related to the way they are curated.

### Correlation between correctly tagged sentences and classification results

We used a collection of 20 manually annotated documents at the sentence level to assess the performance of the string-matching method based on protein features to tag evidence sentences. When measuring the impact on the classifier’s performance, results show that there is no correlation between correctly tagged sentences and correctly predicted categories. We have two hypotheses for this lack of correlation. First, we believe that just splitting the sentences that contain accession information would be sufficient for the classifier to more effectively learn the category features. Even if the sentences did not have exactly the evidence used by biocurators, the most important would be to recall sentences with protein information. Indeed, while the correlation of the classifier’s precision with the tagging method precision is only 0.02, the correlation between the classifier’s precision and the number of positive tagged sentences is 0.26. Notice that there is only a marginal improvement from the *not tagged* to *tagged* model if we consider micro average F1-score (0.71 to 0.72). Thus, the recall increase in positive sentences would provide the edge for the *tagged* model. The second and straightforward hypothesis is that the lack of correlation is related to the small size of the annotated set. To investigate both assumptions, a larger annotated set would be needed, and it is out of the scope of this paper.

### Limitations

Abstract composes the majority of the collection used in the experiments. Nevertheless, a large body of the annotation evidence is expected to be found in the methods and results sections of full text articles. The logistic regression and SVM methods used the whole document in the classification; however, compressed in a 200-vector, which were the best value found during the training phase. On the other hand, for the CNN models the documents were truncated in 1500 tokens (1500 for each branch in the tagged model) due to performance reasons, which limit they comparison with the baselines.

In UniProtKB, a single article can be annotated for many proteins, sometimes with the same class. Hence, the prediction results are not independent, biasing the assessment. We tried to mitigate this issue by creating a test set with unique document à categories pairs; however, examples in this set still cannot be considered as fully independent, as one document can be associated to class sub-sets, e.g., PMID à [Function] and PMID à [Function, Sequences].

## Conclusion

To provide a systematic way of categorizing computationally mapped publications in UniProtKB, in this paper we investigated the use of a supervised CNN classifier for assigning categories to pairs of document-protein accessions. To overcome the issue of multiple category sets associated to a single publication, we proposed an effective strategy to tag publication sentences with protein features and create different feature sets out of a single document entry, which are then fed to different CNN layers. Results showed statistically significant improvements upon models that use only the publication as classification features, improving F1-score up to 22% and 12% when compared to a logistic regression baseline for the micro and macro averages, respectively. Moreover, our results show that text classification supported by CNN models provided an effective way for classifying publications according to the UniProtKB entry categories. As future work, we will investigate whether the classification performance for underperforming classes can be improved by adding expert knowledge into the model. Furthermore, we want to explore whether sentence used as evidence for categorisation can be relocated automatically from the classifier model.

## Funding

ELIXIR-EXCELERATE is funded by the European Commission within the Research Infrastructures programme of Horizon 2020, grant agreement number 676559.

## Conflict of interest

None declared.

